# Adaptive Reconstruction of Pupil Dynamics During Blink Data Loss: A Physiologically-Informed Exponential Recovery and Noise Modelling Framework

**DOI:** 10.1101/2025.08.18.670842

**Authors:** Mohammad Ahsan Khodami, Seyed Mohammad Hosseini, Sofia Kireeva

## Abstract

This paper presents a novel, physiologically inspired framework for reconstructing missing pupil data during blink-induced interruptions. Pupillometry has become integral to studies of cognition, emotion, and neural function, yet the frequent data gaps caused by blinks can compromise the accuracy of inferences drawn from pupil measurements. This approach addresses these challenges by introducing an exponential recovery model that dynamically adjusts its time constant in proportion to blink duration, thus capturing the non-linear dynamics of pupil responses more effectively than traditional interpolation methods. A localised noise estimation procedure further enhances realism by incorporating both signal-dependent and baseline noise derived from statistical properties of the data surrounding each blink. Additionally, a Savitzky–Golay filter is selectively applied to the reconstructed intervals, preserving key physiological features while mitigating high-frequency artefacts. To facilitate the implementation of our framework, we introduce PRPIP, a Python package that implements our methodology. This work demonstrates the effectiveness of this method using empirical pupil data from visual tasks, illustrating that the reconstructed signals maintain physiological fidelity across a range of blink durations. These findings underscore the potential of combining physiological principles with data-driven noise modelling to generate robust, continuous pupil traces. By offering a more accurate reconstruction of pupil size, this framework stands to advance pupillometric research in fields ranging from cognitive psychology to clinical neuroscience.

## Introduction

The pupillometry reveals a complex interplay between physiological mechanisms and cognitive processes ^1^, extending well beyond the pupil’s primary function of regulating light entry into the retina. As a central feature of the human eye ^2,3^, the pupil adjusts its size to control the amount of light that reaches the retina in response to both environmental ^4^ and cognitive stimuli^5^. However, the pupil’s significance goes far beyond its fundamental role in vision; it has emerged as a valuable marker in neuroscience and psychology ^2,6–8^, providing insights into cognitive processes ^7^, emotional states ^9^, and various neurological conditions ^2^.

Pupil size indicates autonomic nervous system activity ^10^, reflecting the interplay between sympathetic and parasympathetic pathways ^11^. The dynamics of pupil size are frequently utilised to investigate states of arousal ^12^, attentional focus ^13–15^, and the decision-making process. Specifically, pupillary responses show characteristic patterns of constriction and dilation that follow physiologically based exponential recovery curves, reflecting underlying smooth muscle activity. These complex dynamics give researchers a valuable, non-invasive perspective on intricate cognitive and physiological phenomena. Consequently, precise pupil size measurement has become an essential tool for scholars in various disciplines.

Modern eye-tracking technologies have transformed our ability to capture and analyse pupillometric responses accurately. These technologies are essential for recording task-evoked changes in pupil diameter in real-time, facilitating the investigation of pupil dynamics across various experimental paradigms. The raw pupillary data collected by eye-tracking devices reflect the physical size of the pupil ^16^, which is typically pre-processed into a standardised format for analysis ^17^. Following data acquisition, rigorous preprocessing becomes essential to extract meaningful and reliable insights from the raw pupillary signals.

The preprocessing pipeline typically involves filtering to eliminate artefacts and noise, smoothing valid samples, and applying baseline corrections. However, the seemingly straightforward task of measuring pupil size is accompanied by complex methodological challenges (for a comprehensive review, see Palegatti et al, (2024) ^18^). Despite these preprocessing efforts, eye-tracking data often faces interruptions from factors such as blinks, head movements, and system noise, leading to gaps, or “pupil loss,” in the pupil measuring time series ^16,19^. This highlights the critical importance of advanced reconstruction techniques to maintain data continuity and reliability. The integrity of pupillometric data is paramount for drawing valid scientific conclusions about underlying cognitive and physiological processes.

Recovering missing pupil data poses a significant methodological challenge that demands advanced scientific approaches. This reconstruction is vital because pupil signals exhibit continuity over time. Pupil dynamics typically evolve gradually over hundreds of milliseconds to seconds, influenced by the smooth muscle activity responsible for dilation and constriction ^20^. Overlaying these slow variations are subtle, rapid fluctuations resulting from fine-grained muscle activity or measurement noise ^21^.

Numerous models have been developed to reconstruct missing pupil data to address the associated challenges. Basic methods, such as linear interpolation ^22^, offer a straightforward solution for bridging small gaps; however, they often fail to capture the complexities of pupil dynamics, particularly during extended interruptions. Spline-based techniques, including cubic ^23^ and Akima splines ^24^, yield smoother reconstructions but may introduce artefacts, such as overshooting, especially at the edges of blink intervals. More sophisticated computational approaches, particularly machine learning techniques ^25^, have emerged as promising alternatives. Nevertheless, these approaches frequently struggle to preserve the intrinsic physiological characteristics of pupil dynamics ^26–28^. Current reconstruction methods often rely on fixed parameters that fail to adapt to varying blink durations ^29–31^, create arbitrary thresholds between different techniques ^26^, and produce unrealistically smooth signals ^17^ that lack the natural variability inherent in physiological data. Additionally, many methods fail to capture the inherently exponential nature of pupillary recovery processes ^32–34,34,35^.

Given these methodological challenges in pupil data reconstruction, this study introduces a novel exponential recovery model that addresses key limitations of existing approaches. the model incorporates dynamic time constant adjustment based on blink duration, unified reconstruction mechanisms across all gap lengths, and physiologically realistic noise characteristics. By combining theoretical insights from pupillary dynamics with practical computational considerations, this work aims to develop a reconstruction method that preserves the intrinsic temporal and physiological characteristics of pupil signals while maintaining computational efficiency. This approach balances biological plausibility and practical implementation for pupillometric research across diverse experimental paradigms.

The current model is derived from principles in physiology, psychology, and computational modelling of pupil dynamics. It builds upon the exponential recovery behaviour of the pupil, a phenomenon characterised by rapid initial constriction followed by a gradual return to baseline^26^. This behaviour is mathematically represented through an exponential recovery function, capturing the non-linear dynamics observed in empirical studies ^6,20,36^. These exponential models are widely supported by empirical pupillometry research, as they accurately represent the non-linear return to baseline following blink events. The model further incorporates a dynamic adjustment of the recovery time constant Tau (τ), which adapts to the duration of the blink. This adjustment reflects the biological reality that longer blinks demand slower recovery processes, while shorter disruptions recover more rapidly, aligning with theoretical insights into physiological variability ^21^.

Additionally, incorporating Gaussian noise, aligned to the study by ^37^ and studies like ^37–43^, scaled to the recovery magnitude, mirrors the natural variability inherent in pupillary responses. The model ensures compatibility with observed data by accounting for signal-dependent and independent noise components, enhancing its realism ^44–46^. The unified approach used in this model applies a single recovery mechanism regardless of blink duration, avoids arbitrary thresholds, and reflects the continuity of underlying physiological processes. Together, these elements demonstrate the model’s firm grounding in theoretical principles while maintaining practical applicability to experimental contexts ^47,48^.

A locality-based approach to noise estimation further enhances the current model. Rather than applying uniform noise parameters across all reconstructed segments, the model evaluates local statistical properties of the pupil signal surrounding each blink event. This localised estimation captures the temporal variability in pupil dynamics that may change throughout an experimental session due to factors such as fatigue, arousal fluctuations, or measurement conditions. By computing the ratio between the local standard deviation and range of the pupil signal within windows surrounding individual blinks, the model derives a noise scale parameter that adaptively reflects the signal’s inherent variability. This approach acknowledges that physiological systems exhibit non-stationary behaviour, with noise characteristics that may evolve over time rather than remain constant throughout a recording session ^17,28^.

Additionally, the model incorporates targeted signal processing through the application of a Savitzky-Golay filter exclusively to the reconstructed blink segments. Unlike global smoothing approaches that may over-attenuate meaningful physiological fluctuations, this selective filtering preserves the natural dynamics of the recorded signal while removing reconstruction artefacts. The Savitzky-Golay filter, with its polynomial fitting approach, maintains critical signal features such as peaks and valleys while effectively reducing high-frequency noise ^49,50^. This targeted filtering strategy strikes a balance between signal fidelity and noise reduction, ensuring that the reconstructed segments integrate seamlessly with the surrounding recorded data. By combining locality-based noise estimation with selective polynomial smoothing, the model produces reconstructed pupil signals that maintain both the statistical properties and physiological characteristics of the original data, facilitating more reliable inferences in pupillometric research.

### Methodology

Research has demonstrated that the pupil’s response to light and subsequent recovery follows an exponential decay pattern ^32,51^. This behaviour, characterised by a rapid initial change followed by a gradual return to baseline, has been further supported by recent evaluations of pupillary light response models ^47,52,53^. Exponential models have been shown to perform comparably to linear models regarding predictive accuracy while better capturing the inherent nonlinear dynamics under varying light conditions ^54–56^. The main formula to reconstruct the pupil during the recovery phase following a blink is defined by

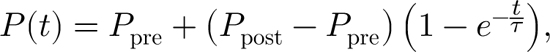

where *P* (*t*) denotes the reconstructed pupil size at time *t*, *P*_pre_ is the pupil size immediately before the blink, *P*_post_ is the pupil size immediately after the blink, and *t* represents the elapsed time during the blink. In our implementation, which is based on measurements in milliseconds, the effective recovery time constant τ is dynamically adjusted according to the blink duration. Specifically, we define

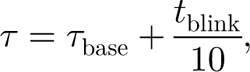

where τ_base_ represents the inherent recovery speed of the pupil and *t*_blink_ is the blink duration. The factor 10 is chosen such that each additional 10 ms of blink time increases τ by 1 unit, a scaling that has been empirically validated to yield physiologically realistic recovery curves. For short blink durations, the effective time constant remains close to τ_base_, whereas for longer blinks, the recovery is proportionally slowed.

To facilitate parameter estimation from experimental data, the pupil size is recorded at various time points *t*_*i*_ during the recovery phase, yielding corresponding measurements *P*_*i*_. To simplify the fitting process, the data is normalised by defining

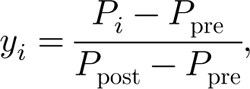

which transforms the model into

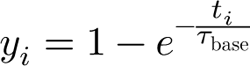

By rearranging this expression as

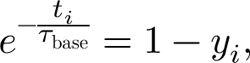

and taking the natural logarithm, we obtain

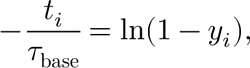

from which an individual estimate of the baseline recovery time constant can be derived as

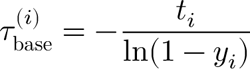

A reliable estimate τ_base_ is then obtained by averaging these individual estimates or by solving the least-squares optimisation problem

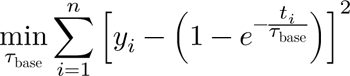

To account for natural physiological variability and measurement uncertainty, the model incorporates a noise component that consists of both signal-dependent and signal-independent elements ^57–59^. The signal-dependent noise variance is modelled by first defining

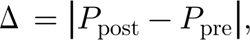

Which represents the magnitude of the change in pupil size. The signal-dependent noise variance is then given by

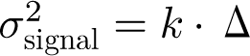

where *k* is a proportionality constant that represents the slope of the linear relationship between the noise variance and the magnitude of the pupil change. When combined with a signal-independent baseline noise variance σ^2^_0_, the total noise variance is expressed as

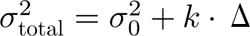

and the overall noise standard deviation becomes

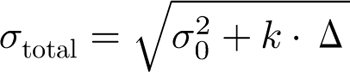

The final reconstructed pupil signal is obtained by adding a Gaussian noise term to the deterministic recovery function ^60–62^, leading to

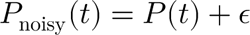

with

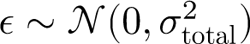

Here, ε represents the Gaussian noise that captures both the inherent physiological variability and the measurement uncertainty, ensuring that the reconstructed pupil data exhibits realistic fluctuations.

In this implementation, the local noise scale *k* is computed by analysing segments of the pupil data immediately surrounding each detected blink event. For each blink, the model defines a local window extending a fixed number of samples *W* before the blink onset and after the blink offset. Within this window, the model computes the rolling standard deviation, denoted as σ_local_, and the rolling range *R*_local_, which is defined as the difference between the maximum and minimum pupil sizes in that window. Mathematically, for a window *W* centred at time *t*, these quantities are given by

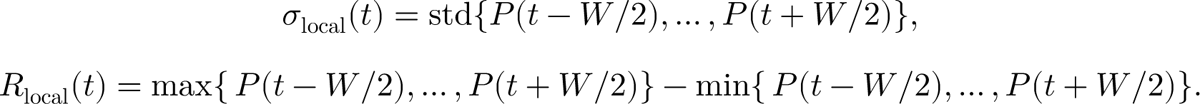

The local noise scale for that blink is then estimated as the ratio

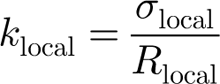

The overall noise scale *k* used in the reconstruction is obtained as the median of these local estimates across all detected blink events, i.e.,

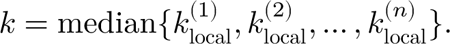

Additionally, a Savitzky–Golay filter ^49,50^ is applied exclusively to the reconstructed segments corresponding to blink intervals to address any irregular artefacts in the reconstructed pupil signal. The Savitzky–Golay filter fits local polynomials ^63–65^ to the data within a moving window; the smoothed value *P*^ (*t*) is computed as

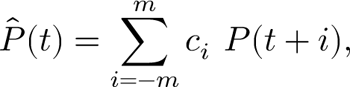

where 2*m* + 1 is the window length, and the coefficients *c*_*i*_ are determined by fitting a polynomial of a specified order to the data within the window. This filtering process effectively reduces high-frequency noise while preserving the pupil response’s overall shape and important transitions.

By employing an exponential recovery function with a dynamically adjusted recovery time constant, a derived noise model based on the data’s local variability, and a targeted smoothing filter for the blink intervals, our approach offers a robust and physiologically grounded method for reconstructing pupil dynamics during blink-induced interruptions.

### Two examples of the method and testing the Model

To evaluate the performance of our proposed reconstruction method, we conducted tests using two distinct examples. Data were recorded during a visual task employing an EyeLink Eye-tracker 1000 Plus Tower setup (SR-Research) with a sample rate of 1000 Hz. Participants were stabilised with a chin rest, and visual stimuli were presented on a monitor with a resolution of 2560×1140 pixels and a refresh rate of 240 Hz. The recordings were performed in a controlled, dim environment, with the participant positioned 57 cm away from the screen, and only the right eye was recorded. The EDF file was converted to CSV format using the etformat package ^66^, then blink was detected by the model provided by Khodami (2025) ^67^ with all analyses conducted in a Python environment.

Trial 1 lasted for a total duration of 6487 milliseconds. Blink detection algorithms identified three distinct blink events, with onsets at 1621, 2508, and 4677 milliseconds and corresponding offsets at 1844, 4517, and 5437 milliseconds. For each blink, a local segment was defined to capture the pupil data surrounding the blink interval. In the first blink, the local segment extended from index 1571 to 1894, and the computed local noise scale was 0.3020. For the second blink, the segment ranged from index 2458 to 4567, yielding a noise scale of about 0.3824, while the third blink, with a segment spanning from index 4627 to 5487, produced a noise scale of approximately 0.3011. The overall average local noise scale, determined as the median of the individual values, was computed to be 0.3020. Subsequently, a Savitzky–Golay filter with a window was applied to each blink segment to smooth the reconstructed signal. Figure 1 shows the result and plot of this process.

**Figure 1.**
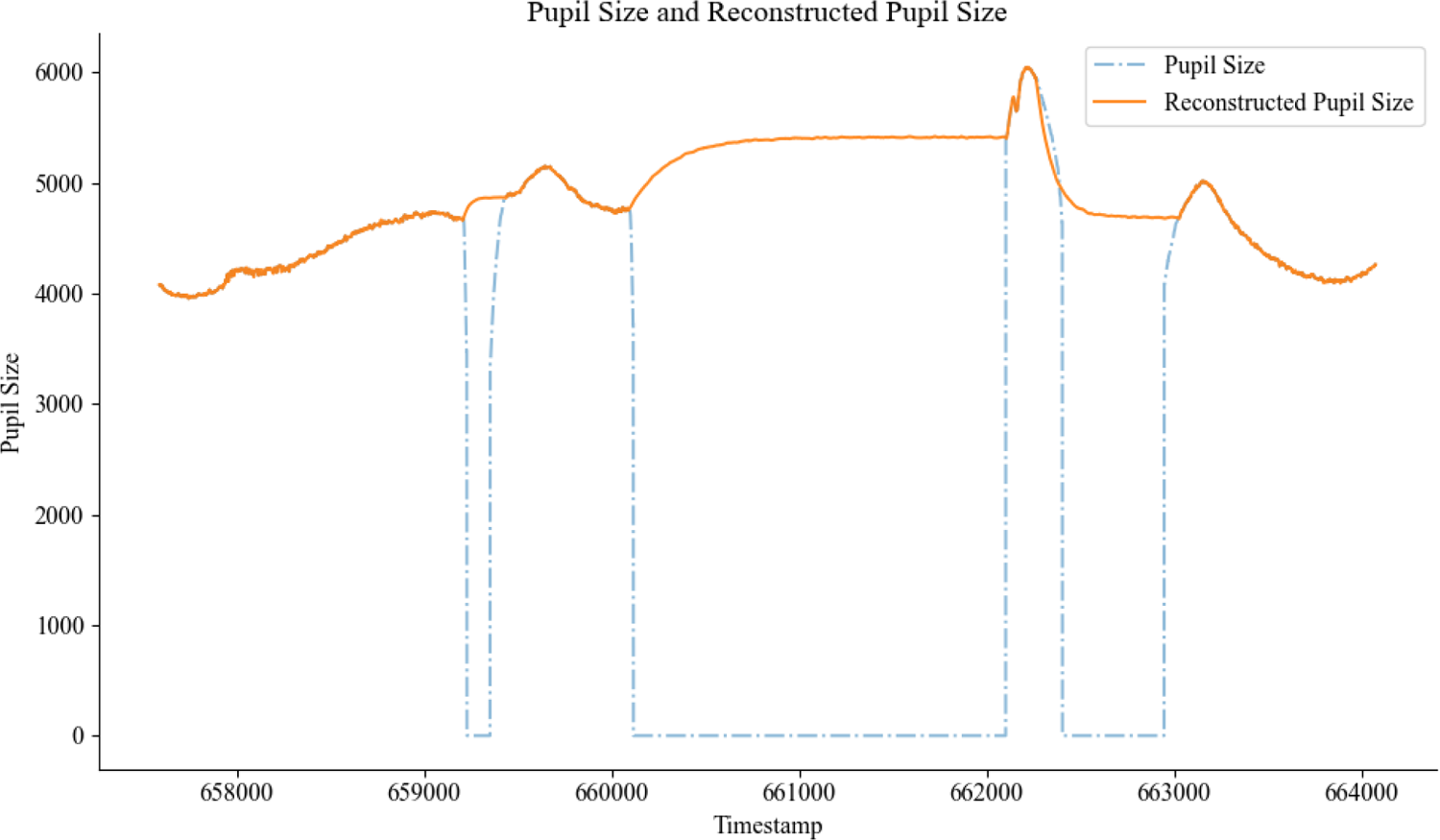
Example of reconstruction using the model. The duration of the total trial is 6847 ms, where we identified 3 blinks at onsets of 1621, 2508, and 4667 and offsets of 1844, 4517, and 5437. After applying the model, the local noise for blinks in order was 0.3019, 0.3823, and 0.3011 and the average local noise of 0.3019 was also computed, and this local noise was applied to reconstruct pupil size. Filter was applied with the window length of 50

**Figure 2.**
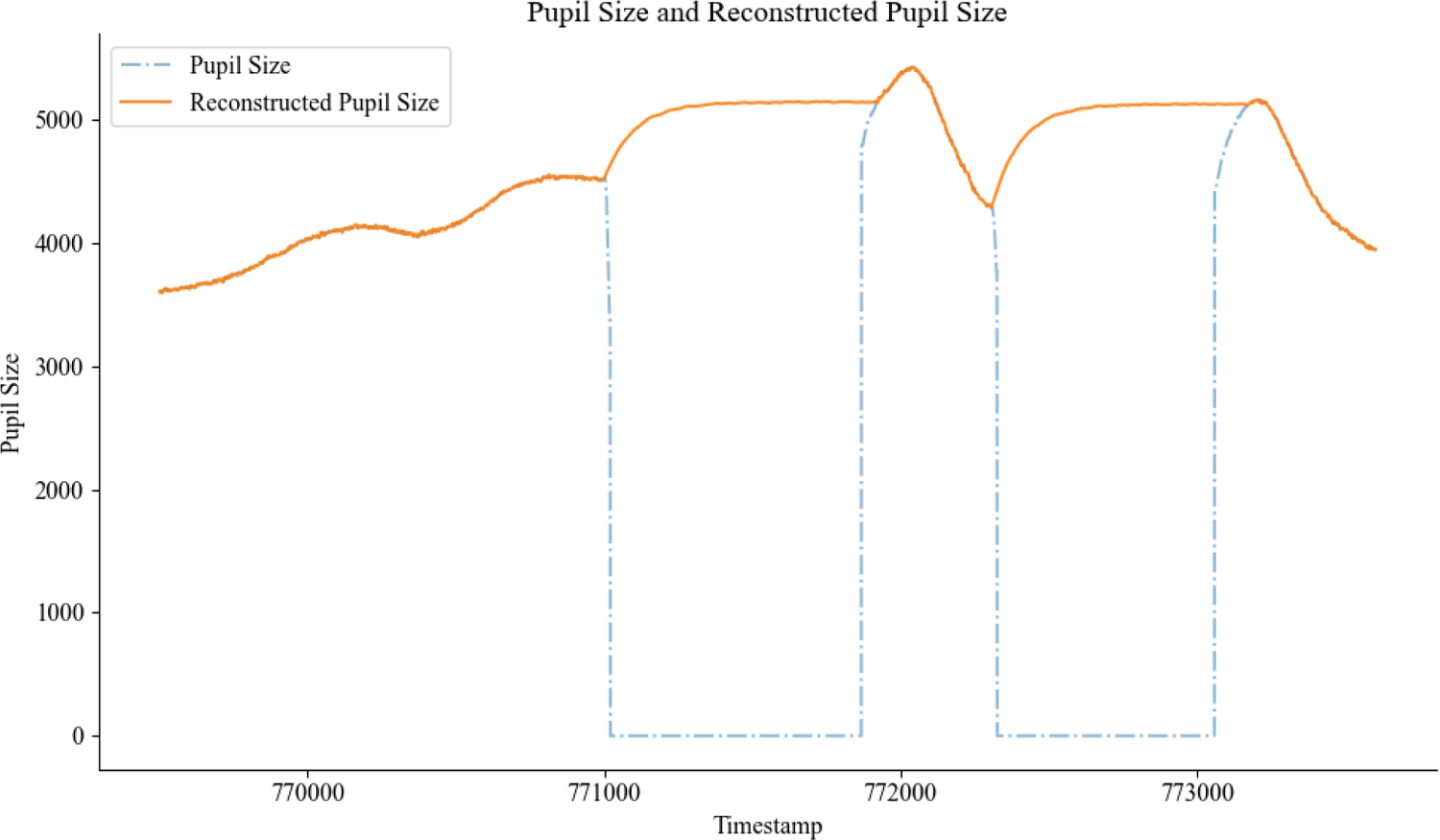
Example of reconstruction using the model, the duration of the total trial is 4103 ms, where we identified 2 blinks at onsets of 1501 and 2420 and offsets of 2808 and 3672, respectively, considering the start of the trial as 0. After applying the model, the local noise for blinks in order was 0.3108 for both and the average local noise of 0.3063, as well as the computed local noise, was applied to reconstruct pupil size.

Trial 2 had a total duration of 4103 milliseconds. Identified with two blink events, with onsets at 1501 and 2808 milliseconds and corresponding offsets at 2420 and 3672 milliseconds. Each blink was analysed within a localised segment to ensure accurate reconstruction. The first blink, spanning the segment from index 1451 to 2470, exhibited a computed local noise scale of approximately 0.3108. The second blink, analysed within the segment ranging from index 2758 to 3722, had a local noise scale of 0.3018. The average local noise scale, determined as the median of the individual estimates, was computed to be 0.3063. A Savitzky–Golay filter was applied on segments with a window length of 50.

### Advantages of the Model

The current model offers several advantages over existing methods for reconstructing pupil size during blink intervals. First, compared to models that employ fixed recovery time constants τ, our approach dynamically adjusts τ based on the duration of the blink. This adaptation ensures that recovery curves accurately reflect the physiological reality of slower recoveries for longer blinks and faster recoveries for shorter blinks, taken the conclusion from ^68,69^. Fixed-τ models often fail to capture this variability, leading to over-smoothing or abrupt transitions that deviate from the natural dynamics of pupil recovery ^14,69^. The dynamic adjustment of τ provides a physiologically plausible scaling that respects the nonlinear relationship between blink duration and recovery time.

Second, our model employs an exponential recovery function for all blink durations, unlike many existing approaches that rely on piecewise strategies with distinct methods for short and long blinks. This design eliminates arbitrary thresholds, which can create inconsistencies and discontinuities in the reconstructed signal. By applying a single recovery mechanism, the model ensures seamless reconstruction consistent with the continuous nature of physiological processes. This unified approach also simplifies implementation and parameter tuning, as researchers need not determine separate algorithms or thresholds for different blink categories.

Third, the model incorporates Gaussian noise scaled to the recovery magnitude, enhancing realism by mimicking the natural variability observed in pupil size measurements, as research indicates that while Gaussian models are useful ^70^. This feature addresses the shortcomings of more straightforward interpolation-based methods that often yield overly smooth reconstructions, lacking the subtle fluctuations seen in empirical data ^71,72^. The reconstructed signal aligns closely with real-world observations by explicitly modelling noise dynamics ^70,73^, maintaining signal-dependent and independent variability. Our approach to estimating the noise scale *k* from local data segments surrounding each blink ensures that the noise characteristics are tailored to the specific recording conditions and individual differences in pupil variability, providing a more accurate representation than models using fixed noise parameters [for more detail, read Appendix A.]

Fourth, the physiological validity of our model represents a significant strength. The exponential recovery curve captures the exponential nature of pupil response as described in experimental and theoretical studies ^37,44,46^. In contrast to polynomial-based methods, which are computationally simple but fail to replicate non-linear recovery dynamics ^74,75^, our model aligns with established findings on exponential decay. This ensures a biologically plausible recovery that bridges the gap between computational accuracy and physiological fidelity. The exponential model’s ability to account for the characteristic rapid initial change followed by a gradual asymptotic approach to the post-blink pupil size is vital for accurately capturing the early phases of pupil recovery, which are often critical in cognitive and perceptual studies.

Fifth, the robustness and simplicity of our implementation make it adaptable to a wide range of experimental paradigms. Unlike machine-learning-based approaches, which often require extensive training data and are constrained to specific tasks ^25,76,77^, our model generalises across diverse conditions involving varying blink characteristics, luminance levels, and noise profiles. This flexibility ensures that our approach is both practical and effective, enabling its application to cognitive and perceptual studies where accurate modelling of pupil dynamics is essential. The model’s minimal parameter set—primarily τ_base_, *k*, and σ_0_—facilitates straightforward adaptation to different experimental conditions without the need for extensive retraining or recalibration.

Sixth, our localised estimation of noise parameters provides a significant advantage over global estimation methods. By analysing segments of pupil data surrounding each blink event, our approach captures local variations in signal quality and physiological state that may affect the noise characteristics. This localisation allows for more accurate noise modelling compared to methods that apply a single noise parameter across the entire recording, which may fail to account for temporal variations in signal quality or physiological state (More at Appendix A).

Seventh, the strategic application of the Savitzky-Golay filter exclusively to the reconstructed blink segments represents a targeted approach to smoothing that preserves the integrity of the original signal where it is valid. This selective filtering contrasts with methods that apply global smoothing, which may inadvertently distort important physiological features outside the blink intervals. Our approach maintains the temporal resolution and dynamic range of the original signal while addressing potential artefacts introduced during reconstruction.

Finally, the model integrates dynamic recovery with advanced noise representation to balance computational efficiency and realism. By avoiding the pitfalls of over-complication and maintaining a firm grounding in physiological principles, it provides a reliable and theoretically sound tool for reconstructing pupil size during blink-induced disruptions, making it broadly applicable across experimental contexts. The combination of physiological validity, computational efficiency, and adaptability positions our model as a valuable contribution to the field of pupillometry, addressing key limitations of existing approaches while maintaining practical utility for researchers studying pupil dynamics in diverse experimental settings.

### Prpip - A package in Python for reconstructing pupil

To facilitate the seamless processing of pupil size data and automate the reconstruction methodology described in this work, we have developed a Python package named prpip. This package integrates all key steps, ensuring a streamlined and reproducible workflow. The package supports multiple file formats, allowing researchers to load raw pupillometry data in various standard structures. Blink detection is implemented using the method offered by Khodami (2025)^67^ to refine onset and offset estimation, improving the accuracy of missing data identification. By consolidating all aspects of blink-affected pupillometry data processing into a single, user-friendly package, *prpip* provides a comprehensive solution for researchers analysing pupil dynamics. This tool significantly enhances the efficiency and reproducibility of pupil data analysis, making it accessible for diverse applications in cognitive science, neuroscience, and psychology.

To install it you may use pip command by running following command as shown in Figure 3.

**Figure 3.**
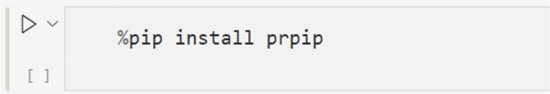
pip install command of installing the prpip package from https://pypi.org. All details and the Wiki are provided at https://pypi.org/prpip (accessed 01 August 2025)

## Discussion

The methodology presented in this paper addresses a significant challenge in pupillometry: the accurate reconstruction of pupil size dynamics during blink intervals. By implementing an exponential recovery model with dynamically adjusted time constants and incorporating both signal-dependent and baseline noise components, our approach successfully generates physiologically plausible pupil size trajectories during periods of data loss.

The primary strength of our method lies in its strong theoretical foundation. The exponential recovery model is congruent with established research on pupillary light response dynamics ^32,51,52,78^. Unlike linear interpolation techniques, which disregard the inherent nonlinear characteristics of pupil recovery, our exponential model accurately captures the rapid initial changes and gradual returns to baseline that characterise actual pupil behaviour ^54,55^.

Additionally, our approach to estimating the noise scale *k* directly from the data represents a significant advancement. By analysing local segments surrounding each blink event and computing the ratio of standard deviation to range, we derive noise parameters that reflect the actual variability in the recorded pupil signals. This data-driven approach ensures that the reconstructed segments exhibit realistic fluctuations that match the surrounding signal characteristics. The adaptive time constant formulation provides another advantage, as it accommodates varying blink durations. This feature is particularly important when processing extended recordings where blink durations may vary considerably due to fatigue or attentional factors.

Finally, the selective application of Savitzky-Golay filtering to the reconstructed segments effectively reduces high-frequency artefacts while preserving the underlying shape of the pupil response curves. This targeted smoothing approach prevents over-filtering of the original recorded data while ensuring seamless transitions between recorded and reconstructed portions of the signal.

### Limitations and Considerations

Despite its strengths, our method has several limitations that warrant consideration. First, the exponential recovery model assumes that the pupil’s behaviour during blinks follows the same dynamics as responses to light stimuli. While theoretically justified, this assumption may not hold under all experimental conditions, particularly during complex cognitive tasks that induce significant pupillary fluctuations.

## Conclusion

This paper presented a comprehensive approach to pupil size reconstruction during blink periods, combining an exponential recovery model with data-driven noise estimation and targeted signal filtering. Our method addresses a critical challenge in pupillometry research and provides a robust solution for generating continuous pupil size traces from recordings affected by blinks.

The implementation’s key features include: (1) a physiologically grounded exponential recovery function, (2) dynamic adjustment of recovery time constants based on blink duration, (3) local estimation of noise parameters from surrounding data segments, and (4) selective application of Savitzky-Golay filtering to ensure smooth transitions between recorded and reconstructed data.

By advancing the methodology for pupil data preprocessing, this work contributes to the broader goal of establishing pupillometry as a reliable tool for investigating cognitive processes, emotional responses, and neurological conditions. The ability to maintain signal continuity despite blink-related interruptions represents an important step toward fully harnessing pupillary measures’ potential as non-invasive windows into neural activity.

## Declarations

### Funding

MAK has received funding from the European Union’s Horizon 2020 research and innovation programme under the Marie Skłodowska-Curie grant agreement No 101034319 and from the European Union – NextGenerationEU; SMH is under doctoral Student fund from University of London, Bayes Business School. SK declare no fundings.

### Conflict of interest

The authors declare that there are no conflicts of interest.

### Ethics approval Not Applicable

Consent for participate Not Applicable

### Consent for publication

All authors affirm their agreement to the publication of this work.

### Availability of data and materials

Both Trial 1 and 2 Data and plotting code is available at https://osf.io/gsr7q/ Code availability The Python package PRPIP, designed for reconstructing pupil data, is available for download and installation from the Python Package Index (PyPI). Installation can be executed via the command provided therein or by adhering to the instructions available at https://pypi.org/project/prpip/. Additionally, plotting code can be obtained from https://osf.io/gsr7q/.

### Author contribution

MAK: Responsible for the development of the methodology, formulation of mathematical concepts, coding, data analysis, writing, editing, drafting, and conceptualisation. SMH: mathematical concepts, code editing, editing. SK: Mathematics, Code, Review

## Acknowledgements

The authors would like to thank Professor Jeremy Wolfe and the Visual Attention Lab (VAL) at Harvard Medical School, along with all VAL Lab members, for their constructive feedback and suggestions. They also appreciate comments and revisions from colleagues, particularly mathematical feedback, and from Dr. Yash Vyas of the Department of Industrial Engineering, University of Padua.

## Appendix A

### Local Noise Addition for Signal Reconstruction

Signal reconstruction is a critical aspect of physiological and natural signal processing, particularly when dealing with missing data. Our research proposes a local noise addition approach as a method to enhance reconstructed pupil size signals, introducing a physiological feature that better reflects the natural variability of the data. This approach acknowledges the inherent noise present in biological systems and integrates it into the reconstruction process, aiming to improve both realism and fidelity. To substantiate our findings, we conducted a comparative analysis of various reconstruction methods, drawing insights from existing literature and empirical findings.

The distinction between local and global reconstruction approaches has been extensively studied. Local methods, as demonstrated by ^37^, have been particularly effective in reconstructing frequency-modulated signals with substantial missing data. Their approach utilised a sliding window technique combined with an iterative orthogonal matching pursuit algorithm, successfully handling missing data rates of 45% and 50% while preserving structural similarity. This finding aligns with our hypothesis that locally applied transformations, such as noise addition, can enhance the fidelity of reconstructed signals.

Global methods, such as the Bayesian inference-based technique employed by ^79^, have shown remarkable accuracy, achieving an *R*^2^ statistic greater than 0.95 in synthetic graph signal reconstructions. However, such methods typically require extensive computational resources and are highly dependent on structured missing data patterns. In contrast, the Environmental Space Time Improved Compressive Sensing (ESTI-CS) algorithm introduced ^80,81^ demonstrated robustness even in the presence of 90% missing data, maintaining an error ratio of 20% or lower. These results emphasise the strength of global approaches in extreme data loss conditions but do not necessarily account for the stochastic fluctuations observed in physiological signals.

The necessity of local adaptations in signal fidelity preservation has been underscored in multiple domains. As measured ^37^ reconstruction quality using the structural similarity index, a metric that aligns with our objective of ensuring physiological plausibility in pupil size reconstruction. Similarly, methods reviewed ^42^ In proteomics data reconstruction, the importance of retaining signal characteristics specific to the domain was emphasised, particularly in mass spectrometry datasets, where local similarity-based approaches such as REM and LSA proved effective.

Computational efficiency also plays a crucial role in method selection. Local reconstruction techniques tend to be computationally efficient ^82^, as observed in studies which described their local fusion frame-based method as having relatively low complexity. In contrast, Bayesian inference methods and global compressive sensing approaches often involve higher computational costs ^79–81^. Other works demonstrated that the RecPF algorithm reduced per-iteration costs compared to the Two-step Iterative Shrinkage/Thresholding (TwIST) method for MRI reconstruction, reinforcing the trade-off between computational expense and reconstruction accuracy ^34^.

Our proposed method of local noise addition contributes to this discourse by leveraging the strengths of local approaches while incorporating stochastic variability intrinsic to physiological signals. The structural similarity preservation observed supports the feasibility of noise addition as a mechanism for improving pupil size reconstruction without excessive computational demand ^83^. Furthermore, graph-based approaches suggest that localised modifications, particularly those that account for smoothness and local-set dependencies, can be advantageous in reconstructing signals with missing data ^42,84^.

The findings reinforce the significance of local methods in reconstructing signals where physiological variations are integral to the underlying process. By incorporating local noise, we align with principles observed in time-series reconstructions, sensor network data, and proteomics signal recovery. This approach not only enhances the realism of the reconstructed pupil size but also maintains the computational advantages inherent to local techniques. Future work will involve empirical validation of this method across diverse missing data scenarios, further solidifying its role in physiologically plausible signal reconstruction.

## Notes

https://osf.io/gsr7q/

### Competing Interest Statement

The authors have declared no competing interest.

## References

1. Pandey, A., Hardingham, N. & Fox, K. Hebbian and homeostatic plasticity mechanisms are segregated in sub-types of layer 5 neuron in the visual cortex. (2022) doi:10.1101/2022.02.11.480060.

2. Zele, A. J. & Gamlin, P. D. Editorial: The pupil: Behavior, anatomy, physiology and clinical biomarkers. Front. Neurol. 11, (2020).

3. Bremner, F. D. THE PUPIL: ANATOMY, PHYSIOLOGY, and CLINICAL APPLICATIONS: By irene E. Loewenfeld. 1999. Oxford: Butterworth-heinemann. Price pound180. Pp. 2278. ISBN 0-750-67143-2. Brain J. Neurol. 124, 1881–1883 (2001).

4. Luchowski, R. et al. Light-modulated sunscreen mechanism in the retina of the human eye. J. Phys. Chem. B 125, 6090–6102 (2021).

5. Mathôt, S. Tuning the senses: How the pupil shapes vision at the earliest stage. Annu. Rev. Vis. Sci. (2020) doi:10.1146/annurev-vision-030320-062352.

6. Joshi, S. & Gold, J. Pupil size as a window on neural substrates of cognition. Trends Cogn. Sci. 24, 466–480 (2019).

7. Einhäuser, W. The pupil as marker of cognitive processes. in Computational and cognitive neuroscience of vision 141–169 (Springer Singapore, 2016). doi:10.1007/978-981-10-0213-7%5C_7.

8. Laeng, B., Sirois, S. & Gredeback, G. Pupillometry: a window to the preconscious? Perspect. Psychol. Sci. 7, 18–27 (2012).

9. Bradley, M., Miccoli, L., Escrig, M. A. & Lang, P. The pupil as a measure of emotional arousal and autonomic activation. Psychophysiology 45 4, 602–7 (2008).

10. Alshanskaia, E. I., Portnova, G. V., Liaukovich, K. & Martynova, O. V. Pupillometry and autonomic nervous system responses to cognitive load and false feedback: an unsupervised machine learning approach. Front. Neurosci. 18, (2024).

11. Szabadi, E. Functional organization of the sympathetic pathways controlling the pupil: Light-inhibited and light-stimulated pathways. Front. Neurol. 9, (2018).

12. Meissner, S. N. et al. Self-regulating arousal via pupil-based biofeedback. *Nat*. Hum. Behav. 8, 43–62 (2023).

13. Lapborisuth, P., Koorathota, S. & Sajda, P. Pupil-linked arousal modulates network-level EEG signatures of attention reorienting during immersive multitasking. J. Neural Eng. 20, 046043 (2023).

14. Thurman, S., Cohen Hoffing, R., Garcia, J. & Vettel, J. Feature-based attention modulates pupil responses by target similarity in a rapid dynamic attention task. J. Vis. 22, 3387 (2022).

15. Salvaggio, S., Andres, M., Zénon, A. & Masson, N. Pupil size variations reveal covert shifts of attention induced by numbers. Psychon. Bull. Rev. 29, 1844–1853 (2022).

16. Ehlers, J., Strauch, C., Georgi, J. & Huckauf, A. Pupil size changes as an active information channel for biofeedback applications. Appl. Psychophysiol. Biofeedback 41, 331–339 (2016).

17. Mathôt, S. Pupillometry: Psychology, physiology, and function. J. Cogn. 1, (2018).

18. Pelagatti, C., Blini, E. & Vannucci, M. Catching mind wandering with pupillometry: Conceptual and methodological challenges. WIREs Cogn. Sci. (2024) doi:10.1002/wcs.1695.

19. Knapen, T. et al. Cognitive and ocular factors jointly determine pupil responses under equiluminance. PLOS ONE 11, e0155574 (2016).

20. Sobczak, F., Pais-Roldán, P., Takahashi, K. & Yu, X. Decoding the brain state-dependent relationship between pupil dynamics and resting state fMRI signal fluctuation. eLife 10, (2021).

21. Winn, M. B., Wendt, D., Koelewijn, T. & Kuchinsky, S. E. Best Practices and Advice for Using Pupillometry to Measure Listening Effort: An Introduction for Those Who Want to Get Started. Trends Hear. 22, 2331216518800869 (2018).

22. Blu, T., Thévenaz, P. & Unser, M. Linear interpolation revitalized. IEEE Trans. Image Process. 13, 710–719 (2004).

23. Knott, G. D. Interpolating Cubic Splines. vol. 18 (Springer Science & Business Media, 1999).

24. Akima, H. A new method of interpolation and smooth curve fitting based on local procedures. J. ACM JACM 17, 589–602 (1970).

25. Das, W. & Khanna, S. A novel pupillometric-based application for the automated detection of ADHD using machine learning. in Proceedings of the 11th ACM international conference on bioinformatics, computational biology and health informatics 1–6 (ACM, 2020). doi:10.1145/3388440.3412427.

26. Fink, L. et al. From pre-processing to advanced dynamic modeling of pupil data. Behav. Res. Methods 56, 1376–1412 (2023).

27. Pinheiro, J. & Bates, D. Mixed-Effects Models in S and S-PLUS. (Springer science & business media, 2000).

28. Kret, M. E. & Sjak-Shie, E. E. Preprocessing pupil size data: Guidelines and code. Behav. Res. Methods 51, 1336–1342 (2018).

29. Chen, S. & Epps, J. Efficient and robust pupil size and blink estimation from near-field video sequences for human–machine interaction. IEEE Trans. Cybern. 44, 2356–2367 (2014).

30. Liang, R. & Song, Q. Blink detection and duration estimation by using adaptive threshold with considering individual difference. in 2021 IEEE international conference on real-time computing and robotics (RCAR) 1116–1121 (IEEE, 2021). doi:10.1109/rcar52367.2021.9517456.

31. Baptista, M. S., Bohn, C., Kliegl, R., Engbert, R. & Kurths, J. Reconstruction of eye movements during blinks. Chaos Interdiscip. J. Nonlinear Sci. 18, (2008).

32. Hansen, R. M. & Fulton, A. Pupillary changes during dark adaptation in human infants. Invest. Ophthalmol. Vis. Sci. 27, 1726–1729 (1986).

33. Pant, M., Zele, A. J., Feigl, B. & Adhikari, P. Light adaptation characteristics of melanopsin. Vision Res. 188, 126–138 (2021).

34. Fang, L.-Y. & Young, R. S. Fourier components of the pupillary response and their relationship to temporal characteristics of luminance and chromaticity processes. in Vision science and its applications TuA2 (Optica Publishing Group, 1995). doi:10.1364/vsia.1995.tua2.

35. Granda, A. M., Dearworth, J. R., Kittila, C. A. & Boyd, W. D. The pupillary response to light in the turtle. Vis. Neurosci. 12, 1127–1133 (1995).

36. Grujic, N., Polania, R. & Burdakov, D. Neurobehavioral meaning of pupil size. Neuron 112, 3381–3395 (2024).

37. Amir, N., Tishby, N. & Nelken, I. A simple model of the attentional blink and its modulation by mental training. PLOS Comput. Biol. 18, e1010398 (2022).

38. Iacoviello, D. & Lucchetti, M. Parametric characterization of the form of the human pupil from blurred noisy images. Comput. Methods Programs Biomed. 77, 39–48 (2005).

39. Dutta, S., Burk, J., Santer, R., Zwiggelaar, R. & Boongoen, T. Noise profiling forANNs: a bio-inspired approach. in Advances in computational intelligence systems 140–153 (Springer Nature Switzerland, 2024). doi:10.1007/978-3-031-47508-5%5C_12.

40. McGill, W. J. & Teich, M. C. A unique approach to stimulus detection theory in psychophysics based upon the properties of zero-mean Gaussian noise. J. Math. Psychol. 33, 99–108 (1989).

41. Selvam, A. M. Noise or random fluctuations in physical systems: a review. in Self-organized criticality and predictability in atmospheric flows 41–74 (Springer International Publishing, 2017). doi:10.1007/978-3-319-54546-2%5C_2.

42. Webb-Robertson, B.-J. M. et al. Review, Evaluation, and Discussion of the Challenges of Missing Value Imputation for Mass Spectrometry-Based Label-Free Global Proteomics. J. Proteome Res. 14, 1993–2001 (2015).

43. Kassinopoulos, M. & Mitsis, G. D. Physiological noise modeling in fMRI based on the pulsatile component of photoplethysmograph. NeuroImage 242, 118467 (2021).

44. van Kempen, J. et al. Behavioural and neural signatures of perceptual decision-making are modulated by pupil-linked arousal. eLife 8, (2019).

45. Geurts, L. S., Ling, S. & Jehee, J. F. M. Pupil-linked arousal modulates precision of stimulus representation in cortex. J. Neurosci. 44, e1522232024 (2024).

46. Weijs, M. L. et al. Pupil self-regulation modulates markers of cortical excitability and cortical arousal. (2024) doi:10.1101/2024.09.04.611153.

47. van der Wel, P. & van Steenbergen, H. Pupil dilation as an index of effort in cognitive control tasks: A review. Psychon. Bull. Rev. 25, 2005–2015 (2018).

48. Bismark, A. W., Mikhael, T., Mitchell, K., Holden, J. & Granholm, E. Pupillary responses as a biomarker of cognitive effort and the impact of task difficulty on reward processing in schizophrenia. Schizophr. Res. 267, 216–222 (2024).

49. Schafer, R. W. What is a savitzky-golay filter?[lecture notes]. IEEE Signal Process. Mag. 28, 111–117 (2011).

50. Krishnan, S. R. & Seelamantula, C. S. On the selection of optimum Savitzky-Golay filters. IEEE Trans. Signal Process. 61, 380–391 (2012).

51. Gramatikov, B., Irsch, K. & Guyton, D. Optimal timing of retinal scanning during dark adaptation, in the presence of fixation on a target: the role of pupil size dynamics. J. Biomed. Opt. 19, (2014).

52. John, B., Raiturkar, P., Banerjee, A. & Jain, E. An evaluation of pupillary light response models for 2D screens and VR HMDs. in Proceedings of the 24th ACM symposium on virtual reality software and technology 1–11 (ACM, 2018). doi:10.1145/3281505.3281538.

53. David-John, B., Raiturkar, P., Banerjee, A. & Jain, E. An evaluation of pupillary light response models for 2D screens and VR HMDs. Proc. 24th ACM Symp. Virtual Real. Softw. Technol. (2018) doi:10.1145/3281505.3281538.

54. Bilgehan, B. Efficient approximation for linear and non-linear signal representation. Iet Signal Process. 9, 260–266 (2015).

55. Wei, K., Fu, Y., Zheng, Y. & Yang, J. Physics-based noise modeling for extreme low-light photography. IEEE Trans. Pattern Anal. Mach. Intell. 44, 8520–8537 (2021).

56. Chen, Z. Y., Peng, Z. S., Yang, J., Chen, W. Y. & Ou-Yang, Z. M. A mathematical model for describing light-response curves in Nicotiana tabacum L. Photosynthetica 49, (2011).

57. Acito, N., Diani, M. & Corsini, G. Signal-dependent noise modeling and model parameter estimation in hyperspectral images. IEEE Trans. Geosci. Remote Sens. 49, 2957–2971 (2011).

58. Jones, K. E., de Hamilton, A. F. C. & Wolpert, D. M. Sources of signal-dependent noise during isometric force production. J. Neurophysiol. 88, 1533–1544 (2002).

59. Brooks, S. Markov chain Monte Carlo method and its application. J. R. Stat. Soc. Ser. Stat. 47, 69–100 (1998).

60. Lalley, S. P. Beneath the noise, chaos. Ann. Stat. 27, (1999).

61. Lev, N., Peled, R. & Peres, Y. Separating signal from noise. Proc. Lond. Math. Soc. 110, 883–931 (2015).

62. Driscoll, M. F. The signal-noise problem — a solution for the case that signal and noise are Gaussian and independent. J. Appl. Probab. 12, 183–187 (1975).

63. Rajagopalan, S. & Robb, R. A. Image smoothing with Savtizky-Golai filters. in Medical imaging 2003: Visualization, image-guided procedures, and display (ed. Galloway, R. L., Jr.) vol. 5029 773 (SPIE, 2003).

64. Schmid, M., Rath, D. & Diebold, U. Why and how savitzky–golay filters should be replaced. ACS Meas. Sci. Au 2, 185–196 (2022).

65. Menon, S. V. & Seelamantula, C. S. Robust savitzky-golay filters. in 688–693 (IEEE, 2014). doi:10.1109/icdsp.2014.6900752.

66. Khodami, M. A. Etformat: A Package for Conversion and Analysis of EDF EyeTracker Data.

67. Khodami, M. A. A Dynamic Threshold-Based Method for Robust and Accurate Blink Detection in Eye-Tracking Data. (2025) doi:10.1101/2025.04.21.649751.

68. Czajka, A. Iris liveness detection by modeling dynamic pupil features. in Handbook of iris recognition 439–467 (Springer London, 2016). doi:10.1007/978-1-4471-6784-6%5C_19.

69. Vincent, P., Parr, T., Benrimoh, D. & Friston, K. J. With an eye on uncertainty: Modelling pupillary responses to environmental volatility. PLOS Comput. Biol. 15, e1007126 (2019).

70. Mangalam, M., Kelty-Stephen, D. G., Hayano, J., Watanabe, E. & Kiyono, K. Quantifying non-Gaussian intermittent fluctuations in physiology: Multiscale probability density function analysis using the Savitzky-Golay detrending. Phys. Rev. Res. 5, (2023).

71. Rodriguez-Perez, D. & Sanchez-Carnero, N. Multigrid/multiresolution interpolation: Reducing oversmoothing and other sampling effects. Geomatics 2, 236–253 (2022).

72. Musial, J. P., Verstraete, M. M. & Gobron, N. Technical Note: Comparing the effectiveness of recent algorithms to fill and smooth incomplete and noisy time series. *Atmospheric Chem*. Phys. 11, 7905–7923 (2011).

73. Kurtz, V. & Lin, H. Kalman filtering with gaussian processes measurement noise. ArXiv abs/1909.10582, (2019).

74. Schramm, T. & Wein, A. S. Computational barriers to estimation from low-degree polynomials. Ann. Stat. 50, (2022).

75. Farina, M. & Piroddi, L. An iterative algorithm for simulation error based identification of polynomial input–output models using multi-step prediction. Int. J. Control 83, 1442–1456 (2010).

76. Fuhg, J. N., Hamel, C. M., Johnson, K., Jones, R. & Bouklas, N. Modular machine learning-based elastoplasticity: Generalization in the context of limited data. Comput. Methods Appl. Mech. Eng. 407, 115930 (2023).

77. Tsialiamanis, G., Dervilis, N., Wagg, D. J. & Worden, K. Towards a population-informed approach to the definition of data-driven models for structural dynamics. Mech. Syst. Signal Process. 200, 110581 (2023).

78. Van der Maas, H. L. & Molenaar, P. C. Stagewise cognitive development: an application of catastrophe theory. Psychol. Rev. 99, 395 (1992).

79. Antonian, E., Peters, G. W. & Chantler, M. Bayesian reconstruction of Cartesian product graph signals with general patterns of missing data. J. Frankl. Inst. 361, 106805 (2024).

80. Kong, L., Xia, M., Liu, X.-Y., Wu, M.-Y. & Liu, X. Data loss and reconstruction in sensor networks. in 2013 Proceedings IEEE INFOCOM (IEEE, 2013). doi:10.1109/infcom.2013.6566962.

81. Kong, L. et al. Data Loss and Reconstruction in Wireless Sensor Networks. IEEE Trans. Parallel Distrib. Syst. 25, 2818–2828 (2014).

82. Aceska, R., Bouchot, J.-L. & Li, S. Local sparsity and recovery of fusion frame structured signals. Signal Process. 174, 107615 (2020).

83. Amin, M., Dogaru, T. & Jokanovic, B. Reconstruction Of Locally Frequency Sparse Nonstationary Signals From Random Samples. Zenodo (2014) doi:10.5281/ZENODO.44160.

84. Gu, Y. & Wang, X. Local-Set-Based Graph Signal Sampling and Reconstruction. in Signals and Communication Technology 255–292 (Springer International Publishing, 2018). doi:10.1007/978-3-030-03574-7%5C_7.

